# Cirrhosis and Inflammation Regulates CYP3A4 Mediated Chemoresistance in Vascularized Hepatocellular Carcinoma-on-a-chip

**DOI:** 10.1101/2022.05.04.490682

**Authors:** Alican Özkan, Danielle L. Stolley, Erik N. K. Cressman, Matthew McMillin, Thomas E. Yankeelov, Marissa Nichole Rylander

**Affiliations:** Department of Mechanical Engineering, The University of Texas, Austin, TX, 78712, United States; Department of Interventional Radiology, The University of Texas MD Anderson Cancer Center, Houston, TX, 77030. United States; Department of Internal Medicine, The University of Texas at Austin, Dell Medical School; Central Texas Veterans Health Care System, Austin, TX, 78712, United States; Department of Biomedical Engineering, The University of Texas, Austin, TX, 78712, United States; Oden Institute for Computational Engineering and Sciences, The University of Texas, Austin, TX, 78712, United States; Departments of Diagnostic Medicine, The University of Texas, Austin, TX, 78712, United States; Department of Oncology, The University of Texas, Austin, TX, 78712, United States; Livestrong Cancer Institutes, Dell Medical School, The University of Texas, Austin, TX, 78712, United States; Department of Imaging Physics, The University of Texas MD Anderson Cancer Center, Houston, TX, 77030; Wyss Institute for Biologically Inspired Engineering at Harvard University, Boston, MA, 02115, United States

**Keywords:** Microfluidics, Organ-on-a-Chip, Hepatocellular Carcinoma, Drug Metabolism, Inflammation, Cirrhosis, Chemoresistance

## Abstract

Understanding the effects of inflammation and cirrhosis on the regulation of drug metabolism during the progression of hepatocellular carcinoma (HCC) is critical for developing patient-specific treatment strategies. In this work, we created novel three-dimensional vascularized HCC-on-a-chips (HCCoC), composed of HCC, endothelial, stellate, and Kupffer cells tuned to mimic normal or cirrhotic liver stiffness. HCC inflammation was controlled by tuning Kupffer macrophage numbers, and the impact of cytochrome P450-3A4 (CYP3A4) was investigated by culturing HepG2 HCC cells transfected with CYP3A4 to upregulate expression from baseline. This model allowed for the simulation of chemotherapeutic delivery methods such as intravenous injection and transcatheter arterial chemoembolization (TACE). We showed that upregulation of metabolic activity, incorporation of cirrhosis and inflammation, increase vascular permeability due to upregulated inflammatory cytokines leading to significant variability in chemotherapeutic treatment efficacy. Specifically, we show that further modulation of CYP3A4 activity of HCC cells by TACE delivery of doxorubicin provides an additional improvement to treatment response and reduces chemotherapy-associated endothelial porosity increase. The HCCoCs were shown to have utility in uncovering the impact of the tumor microenvironment (TME) during cancer progression on vascular properties, tumor response to therapeutics, and drug delivery strategies.

**Statement of Significance:** Regulation of drug metabolism during the cancer progression of hepatocellular carcinoma (HCC) can be influential to develop personalized treatment strategies. We created novel vascularized hepatocellular carcinoma-chip (HCCoC) composed of tunable collagen and four main liver-specific cell lines to be used as a preclinical tool. In this model, we found cancer evolution states such as inflammation and cirrhosis increases vascular permeability progressively as a result of increased inflammatory cytokines. Furthermore, delivery of doxorubicin only with embolization improved treatment efficacy by decreasing CYP3A4 activity, which can modulate treatment outcome. Overall, we found different disease states can be influential on CYP3A4, thus its targeting can improve HCC treatment outcome.

## 1. Introduction

Hepatocellular carcinoma (HCC) is the most common form of liver cancer and has the second-lowest survival rate at 20% among the other cancer types [1]. Despite the increase in treatment options and ongoing clinical trials, HCC had the highest increase in mortality among all cancer types from 2010 to 2020 [1,2]. HCC is commonly diagnosed in the intermediate to advanced stages of disease and is typically associated with underlying chronic liver disease, complicating treatment options [3,4]. Several systemic and localized drug-based therapies are the current standard of treatment for HCC. However, patient responses are highly variable, emphasizing the need to stratify patients for the most effective treatment [5]. Traditional HCC treatments involve intravenous (IV) administration of chemotherapy drugs such as doxorubicin, although more recently, there have been advances in targeted therapies and immunotherapies [6]. Conventional treatments have had minimal impact on mean survival compared to patients who received no treatment [7]. Over the past two decades, the application of localized delivery of chemotherapeutics through transcatheter arterial chemoembolization (TACE) has shown a significant improvement over traditional IV methods to treat HCC [8]. TACE blocks the arterial blood supply to the tumor using particulate or viscous liquid agents such as degradable starch microspheres, drug-eluting beads, or vehicles such as ethiodized oil [9]. This well-established technique increases the chemotherapeutics’ local dose and residence time in the target area, reduces the supply of nutrients and oxygen to the tumor and simultaneously limits drug exposure and toxicity to the rest of the body [10]. As such, TACE has emerged as the primary treatment method for intermediate state HCC patients, including patients with unresectable multinodular lesions. Recently, disease staging systems created by the Barcelona Clinic Liver Cancer (BCLC), Japanese Integrated Staging (JIS), the Chinese University Prognostic Index (CUPI), and the Hong Kong Liver Cancer (HKLC) have all supported the use of TACE for early-stage HCC in patients that do not respond to conventional IV chemotherapies [11]. While TACE provides several advantages over IV administration, previous studies have demonstrated that TACE still results in a high rate of persistent, viable tumor cells in a significant fraction of patients [12]. In addition, drug-metabolizing enzymes play a critical role in patient-to-patient variability [13], and possibly altering treatment outcomes [14]. Therefore, while a substantial survival benefit can be realized, greater characterization of which underlying tumor microenvironment (TME) conditions affect therapy outcome and their contribution to treatment variability observed in IV and TACE treatment of HCC is needed [15–17].

Recent exploration into contributing TME factors that can alter chemotherapy treatment outcomes has highlighted metabolic enzymes such as the cytochrome P450 (CYP) family, primarily the CYP3A4 subgroup predominantly expressed by hepatic cells [13]. In addition, clinical reports indicate that high CYP3A4 expression in HCC patients can contribute to a poor prognosis [18]. Modifications of the liver TME from underlying chronic liver disease associated with increased inflammation and/or cirrhosis have been shown to regulate CYP3A4 activity in HCC cells [19], however the contribution of each factor is currently unknown. Furthermore, current experimental models of HCC treatment are limited by their inability to tune the microenvironment, replicate the complex architecture of the liver sinusoids, maintain a high number of various liver-resident cells, and control the hemodynamics of the TME [20]. These limitations of current HCC models limit insight into CYP3A4 regulation by the TME and the subsequent effects on chemotherapy responses.

Traditionally, two-dimensional (2D) monolayer *in vitro* studies have been used to estimate optimum chemotherapy dosing and treatment combinations for HCC [21,22]. Bowyer *et al*. showed that HCC treatment efficacy associated with combined delivery of doxorubicin and rapamycin increased in response to upregulation of hypoxia-inducible factor 1 (HIF-1) [21]. Furthermore, Coriat *et al*. demonstrated inhibition of HepG2 and HUVEC growth after exposure to sorafenib [22]. However, studies on HCC dosing with the chemotherapeutics doxorubicin and sorafenib showed that cirrhotic 3D collagen hydrogels significantly impact treatment efficacy compared to traditional 2D monolayer studies [19]. 2D *in vitro* cultures do not recapitulate the cellular complexity of the TME, functional transport barrier of the vessels within the liver, or provide a microenvironment with representative stiffness for various disease states, including cirrhosis, limiting applicability to clinical conditions in drug efficacy estimations.

To address the need to develop a more physiologically representative and tunable model of HCC, vascularized liver-on-a-chip (LoC) systems have emerged as a means to mimic and create complex 3D TME architecture to replicate varying disease states and provide multicellular culture under physiological flow conditions for investigating cell-cell, and cell-matrix interactions. These systems allow the incorporation of hemodynamics, complex cellular composition, and a tunable 3D microenvironment architecture [19,23]. Moreover, these systems have been shown to replicate the human liver’s pharmacokinetics and pharmacodynamics (PK/PD) when designed according to allometric scaling laws [24]. Therefore, LoCs could potentially address the challenges associated with testing HCC treatment methods and optimizing the drug development process by more accurately replicating disease progression states. Several LoC models incorporating all of the critical liver-specific cells, have been developed to evaluate the toxicity of compounds. However, these models do not have any extracellular matrix within the devices. Typically, apical and basal channels were separated with a porous membrane, where hepatocytes and stellate cells were cultured in apical channels, and endothelial, and Kupffer-like macrophages that are from THP-1 derived were cultured in the basal channel. A recent LoC study with human primary hepatocytes cultured with native liver-specific cells using a similar system showed the differential effect of drug toxicity compared to dog and rat hepatocytes [25]. Results of this study proved human, and animal models provide different hepatotoxicity levels to therapeutics. This study demonstrated LoCs could be more representative of patient response to therapeutics with the culture of human cells than the animal models. Similarly, another study used LoCs to assess alcohol-associated liver damage using ethanol and lipopolysaccharide to predict alcohol-associated inflammatory responses in hepatocytes [26]. Finally, an *in vivo*-like liver oxygen gradient was accomplished in a glass-based LoC to show that hypoxia contributes to non-alcoholic fatty liver disease-associated lipid accumulation in primary hepatocytes [27]. Although these systems contain liver-specific cells, these models do not recapitulate HCC-related liver disease. Among the relevant microenvironmental factors, they did not incorporate physiological wall shear stress (WSS) on the endothelial cells (EC), capture liver disease states such as inflammation and cirrhosis, or include tunable biomaterials to replicate stiffness-associated response of the ECM. The mechanistic investigation of HCC cells in LoCs and their therapeutic response has been limited. Lee *et al*. investigated the role of T-cell functions on hepatitis B-related hepatocellular carcinoma cells cultured in collagen within a 3D microfluidic device made out of polydimethylsiloxane (PDMS) with continuous culture media supply [28]. In this study, monocytes in 3D suppressed T cell-mediated cytotoxicity toward hepatocellular carcinoma cells. However, comparable observations were not recapitulated in a 2D system in the same study. Unfortunately, there is a lack of HCC-specific LoC model to test the regulation of vascular properties, solute transport, chemotherapeutic efficacy, or CYP3A4 metabolic activity in response to different disease states, that is needed for further research. Furthermore, previous models were not implemented to simulate multiple drug delivery methods such as IV and TACE and measure tissue response for treatment planning.

The present work presents a vascularized HCC-on-a-chip (HCCoC), containing HCC-derived hepatocytes and three other major liver cells-stellate cells, Kupffer-like macrophages, and ECs-and we evaluate the therapeutic efficacy of doxorubicin delivered under different clinically relevant cirrhotic and inflamed cancer stages for the first time. This new disease model also contains the liver-specific space of Disse and incorporation of physiological flow and associated WSS in the vessel. The HCCoC incorporates native HepG2 HCC cells, as well as the upregulated HepG2-CYP3A4 phenotype cells, within representative ECM to assess the impact of CYP3A4 expression on drug response. The HCCoC also recapitulates physiological vascular transport, TME mechanical properties, and liver functional markers, including albumin secretion. Inflammatory cytokine levels increased as a result of cirrhosis and inflammation, modeled by the increase in collagen concentration and Kupffer cell population, respectively, which are similar to those observed in patients. We observed that vascular permeability and endothelium porosity increased when inflammation and cirrhosis were induced, likely stemming from upregulated inflammatory cytokines and CYP3A4 metabolic activity. In HCCoC, we quantitatively predicted the treatment efficacy of IV and TACE drug delivery methods. Our results demonstrate inflammation, cirrhosis, and embolization result in the regulation of CYP3A4 activity in HCC cells, which also controls the treatment efficacy of doxorubicin. Overall, our results propose that the regulation of CYP3A4 expression could be a major factor in doxorubicin treatment efficacy in HCC.

## 2. Materials and Methods

### 2.1 Cell Culture

Human hepatocellular carcinoma HepG2 (ATCC) and CYP3A4 transfected HepG2 cells were cultured with 1X DMEM supplemented with 10% heat-inactivated fetal bovine serum (FBS, Sigma Aldrich, MO), and 1% Penicillin/Streptomycin (P/S, Invitrogen, CA). HepG2-CYP3A4 phenotype has been previously generated to express 7.14-fold higher CYP3A4 expression compared to HepG2 (Fig. S1), which could be found elsewhere [19,29]. Human hepatic stellate cell LX-2 was cultured in HEPES-buffered Dulbecco’s modified Eagle medium (DMEM) (Lonza BioWhittaker, Verviers, Belgium) supplemented with 4 mmol/L L-glutamine (Lonza), 100 IU/ml Penicillin/100 mg/ml Streptomycin (Lonza), and 10% heat-inactivated FBS. Human monocyte cell THP-1 (ATCC, Manassas, MA) was cultured in RPMI 1640 medium supplemented with 10% heat-inactivated FBS, 1% P/S, and 0.05 mM 2-mercaptoethanol. THP-1 cells were differentiated into unpolarized macrophages, representative of liver-specific Kupffer cells, by adding 200 ng/mL phorbol myristate acetate (PMA, P1585, Sigma Aldrich) in RPMI 1640 medium for 48 hours followed an additional 6 hours in PMA-free RPMI growth medium before harvesting and seeding. Telomerase immortalized microvascular endothelial cells, stably transduced with mKate red fluorescent protein lentivirus (provided as a generous gift from Dr. Shay Soker at the Wake Forest Institute for Regenerative Medicine (Winston-Salem, NC)), were cultured in Endothelial Basal Medium-2 (EBM-2, Lonza, MD) and supplemented with an Endothelial Growth Media-2 (EGM-2) SingleQuots Kit (Lonza, MD). Since liver sinusoid ECs results in higher (7.13 fold higher) than physiological vascular permeability compared to microvasculature ECs [27], we selected to use microvasculature ECs as the source of ECs. Additionally, primary cells usually have a limited lifespan, frequently change their phenotypes, and have large variability over different batches of isolation, we used stable, immortalized but also representable ECs. All cells were cultured in temperature and humidity-controlled incubator (5% CO_2_, 37°C, Thermo Fisher Scientific, Rochester, NY) and were grown until 70% confluence and used within the first eight passages.

### 2.2. Fabrication of Hepatocellular Carcinoma-on-a-chip

The housing of the microfluidic device was fabricated with PDMS (Silgard 184; Dow Chemical, MI). The device within the PDMS housing consists of a vascularized luminal hollow space surrounded by two layers of ECM gel chamber. Devices were fabricated and assembled as we have previously published (Fig. S2) [23,30]. Briefly, the PDMS layer of the device was fabricated via standard soft lithography using an aluminum master mold produced by micromilling to form the housing of the device. Well mixed and desiccated PDMS is mixed with a curing agent at 10:1 ratio, poured into the aluminum mold, and cured at 70°C for 1 hour. Solidified PDMS was removed, and plasma treated with a glass slide at 30W for 3 minutes to bond them permanently. Compared with previous microfluidic systems from our group, one unique characteristic of the present design is the high number of separate vessels parallel to each other (n=3), allowing multiple experiments in parallel.

Devices were UV sterilized overnight before cell culture. To prevent delamination of collagen gel, devices were treated with 1% Polyethylenimine (P3143, Millipore Sigma, Burlington, MA) for 10 minutes, followed by 0.1% Glutaraldehyde (AAA17876AE, Thermo Fisher Scientific, Waltham, MA) treatment for 20 minutes at 37°C. Devices were then washed twice with sterile deionized water. Rat tail type I collagen was used to recapitulate the TME as it is the primary ECM component of the liver [31,32] and allows repeatable tuning of the microenvironmental stiffness based on controlling collagen concentration.[33] As we have previously published, the stock solution of type I collagen was prepared by dissolving excised rat tail tendons in an HCl solution at a pH of 2.0 for 12 hours at 23°C.[33] The solution was then centrifuged at 4°C for 45 minutes at 30000 g, and the supernatant was collected, lyophilized, and stored at -20°C. The lyophilized collagen was mixed with diluted 0.1% glacial acetic acid, maintained at 4°C, and vortexed every 24 hours for three days to create a collagen stock solution. Finally, collagen was centrifuged at 4°C for 10 minutes at 2700 rpm to remove air bubbles. Collagen concentrations of 4 and 7 mg/mL were used to replicate normal and cirrhotic liver stiffness, respectively, which we have previously demonstrated to match native liver Young’s modulus [23,33].

The first concentric layer of the HCCoC representing the hepatic region was prepared with collagen I on ice at 4 or 7 mg/mL (normal or cirrhotic stiffness, respectively) with HCC cells, HepG2, or HepG2-CYP3A4, at 2×10^6^/mL and subsequently neutralized to pH 7.4 with 1X DMEM, 10X DMEM, and 1N NaOH (Fisher Scientific, Pittsburgh, PA.). The solution was then polymerized around a 22G needle at 37°C. After collagen polymerization, the needle was removed to create a hollow lumen. The second concentric layer representing the space of Disse was prepared as above with 2 mg/mL collagen solution with LX-2 stellate cells at 5×10^6^/mL, injected in the hollow lumen, and polymerized around a 27G needle. After removing the 27G needle, ECs and differentiated THP-1 derived macrophages were introduced to the 27G hollow to form the vascular endothelium. As we have previously published, the resulting vasculature underwent flow preconditioning for three days with incremental WSS [23,30]. The number of differentiated THP-1 derived Kupffer macrophages representing liver resident Kupffer cells were tuned to mimic *in vivo* data of hepatocyte to Kupffer ratio in normal and inflamed liver states [34,35]. A summary of the four states implemented in the HCCoCs is shown below in **Table 1**.

**Table 1:**
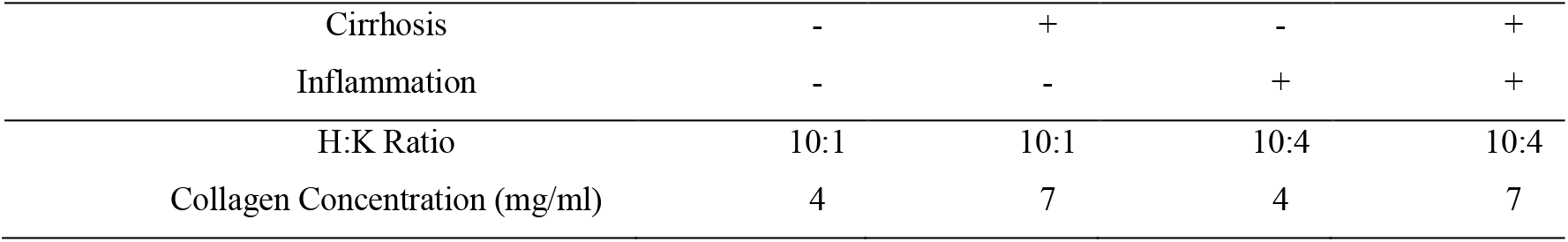
Tested stiffness and disease states in HCCoCs. H:K is HCC:Kupffer ratio.

|  |  |  |  |  |
| --- | --- | --- | --- | --- |
| Cirrhosis | - | + | - | + |
| Inflammation | - | - | + | + |
| H:K Ratio | 10:1 | 10:1 | 10:4 | 10:4 |
| Collagen Concentration (mg/ml) | 4 | 7 | 4 | 7 |

### 2.3. Functional Protein Secretion of the HCC Cells

Albumin, one of the major proteins expressed in the liver, was measured in this study to show the liver cells’ functionality and demonstrate that the model is stable and representative of the liver. Albumin levels were measured using enzyme-linked immunosorbent assay (ELISA). Briefly, after completion of the three-day shear stress conditioning protocol, cell supernatants from the flow outlet were collected after additional exposure to physiological WSS for 6 hours and were stored at -20°C. Albumin secretion was quantified using Human Albumin (ALB) ELISA Kit (EHALB, Invitrogen, CA) according to the manufacturer’s protocol.

### 2.4. Unconfined Compression Test on the HCCoC

The Young’s modulus of collagen hydrogel used for the ECM was quantified to compare the native (4 mg/ml) and cirrhotic (7 mg/mL) states implemented in the HCCoC relative to published clinical data. The Young’s modulus of HCCoC collagen hydrogels was measured with quasi-steady uniaxial unconfined compression. Collagen hydrogels at the same concentration of HCCoC were created in 24 well-plates and isolated with a 1.5 mm diameter biopsy punch. Cylindrical biopsy samples (1.5 mm diameter and 1 mm height) were compressed with 20 mm diameter load cell (10 N Static Load Cell, 2519-10N) attached to an Instron compression machine on a flat steel surface. The load cell was lowered approximately 2 mm away from the base surface and displaced 1.5 mm at a rate of 0.0085 mm/s resulting in 0.1% strain/s over the range of 0-20% strain. Engineering stress was calculated from the force divided by the initial area of the collagen sample. All measurements were performed at room temperature (23°C), and the total duration of each experiment was less than 4 minutes.

### 2.5. Assessment of Vascular Transport Characteristics

Vascular permeability was measured to investigate how the endothelium is altered in response to different TME conditions by measuring the transport of particles of varying solute sizes across the endothelium. Dextran (3 kDa and 70kDa molecular weight) solutions of 10 μg/mL diluted in serum-free endothelial media were perfused for 2 hours at 59 μL/min flow rate, corresponding to physiological 1 dyn/cm^2^ WSS, using a syringe pump (PHD Ultra, Harvard Apparatus, Holliston, MA). The fluorescence intensity was measured with Leica (Wetzlar, Germany) TCS SP8 confocal microscope with a custom enclosure for temperature and atmosphere control (37°C and 5% CO_2_) using 10x (0.6 NA) objective at a resolution of 512 × 512 pixels every 5 minutes [23]. The intensity in the vascular and ECM components was used to calculate vascular permeability (P), under the solute flux *J*_*S*_ acting on per endothelial surface area, which is driven by transendothelial concentration difference Δc:

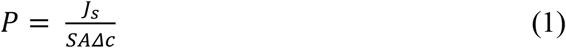

which takes the expanded form:

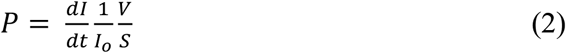

where dI/dt is the intensity change in the ECM over time, *I*_*O*_ is the intensity inside the vessel, V is the volume of the vessel, and S is the vessel’s surface area. Additionally, vascular porosity was quantified using the confocal images of mKate-tagged ECs we have previously published using ImageJ (National Institutes of Health, Bethesda, MD) [23].

### 2.6. Cytokine quantification

17 cytokines secreted from the HCCoC were analyzed using Bio-Plex Pro Human Cytokine Group-I Panel (M5000031YV, BioRad, CA). HCCoCs were exposed to physiological WSS for 6 hours after completion of the preconditioning protocol, and supernatants were collected and stored at -80°C until the measurement.

### 2.7. Drug Treatment and Transport Assessment

The regulation of chemoresistance by TME was investigated by treating vascularized HCCoCs with 10 μM doxorubicin (D1515, Sigma-Aldrich, St. Louis, MO) suspended in endothelial growth media for 24 hours based on standard effective drug concentration established in our previous study [19]. Doxorubicin was either delivered by flowing continuously through the HCCoC vessel to mimic IV or injected with a pipette, and the flow subsequently halted and blocked with sterile pins to mimic embolization associated with TACE.

### 2.8. Post-Treatment Viability Assay

To assess the efficacy of IV and TACE treatments, live and dead cell populations were identified using Calcein and PI staining. After doxorubicin treatment, drug-free media was perfused through the HCCoCs for 24 h. Then, the HCC cell viability in the vascularized HCCoCs was assessed by live/dead staining with Calcein AM (17783, Sigma-Aldrich, St. Louis, MO) and propidium iodide (PI, P1304MP, Invitrogen, Waltham, MA). Fluorescence images were acquired, segmented, and analyzed to detect the number of live and dead cells. Briefly, devices were washed with warm PBS for 15 minutes and treated with 5 μM Calcein AM and 1 μM PI dissolved in warm PBS for two hours before imaging. As an indicator of cell viability, survival fraction was calculated by dividing the number of live cells by the total number of cells.

Nonselective damage to ECs caused by doxorubicin treatment was quantified by measuring vascular porosity of mKate tagged ECs on the lumen using confocal microscopy and image processing in ImageJ. Frangi filter was applied on the projection of 3D confocal images and a fraction of endothelial-free areas compared to the total surface area of the lumen.

### 2.9. Measurement of CYP3A4 Activity Regulation

Hepatocyte metabolic activity, as measured by CYP3A4 expression, has been shown to be highly variable between patients [13] and may serve as a potential marker for HCC chemoresistance. The impact of cirrhosis, inflammation, and drug delivery method (IV and TACE) on CYP3A4 activity was tested with the P450-Glo CYP3A4 assay kit (V9001, Promega, Madison, WI, USA) according to the manufacturer’s protocol. After the three-day preconditioning of the vessel, HCCoC devices underwent continuous perfusion of drug-free endothelial culture media or halting of media flow to serve as controls for IV and TACE procedures, respectively. 24 hours after treatment, the devices containing the cells and ECM were placed in 24 well plates and incubated in culture medium with 3 μM luciferin isopropyl acetal (Luciferin-IPA; a luminogenic substrate for CYP3A4) at 37°C for two hours. The culture media was then transferred to the 96-well opaque white luminometer plate (white polystyrene; Costar, Corning Incorporated), the luciferin detection reagent was added. Luminescence was measured by the Cytation 3 plate reader (BioTek, VT) with fresh culture medium as a blank control.

### 2.10. Statistical Analysis

Each experiment was performed using at least four devices in four biological replicates. Statistical significance was assessed using One-way ANOVA with multiple comparison tests or two-tailed unpaired *t*-test in Matlab (Mathworks, Natick, MA) with p<0.05 regarded as statistically significant. All data are presented as mean ± standard deviation (SD) unless otherwise mentioned.

## 3. Results

### 3.1. Design and Fabrication of Physiologically Relevant HCCoC

As shown in Fig. 1a, the vascularized HCCoC consists of the four major cells in the liver: HCC hepatocyte, stellate, Kupffer, and endothelial cultured in a 3D collagen type 1 extracellular matrix (ECM). The HCCoC with cells and ECM is housed in a gas-permeable PDMS holder. Concentric layers of cells and collagen matrix were utilized to replicate the physiological structure of the liver sinusoid. The aligned vascular endothelium is established by continuously flowing culture medium and increasing the WSS to 1 dyn/cm^2^, which matches the WSS of the human liver TME, over three days according to published protocols (Fig.s 1b and S3) [23]. The central vascular channel consists of preconditioned, confluent, aligned ECs and Kupffer macrophages (Fig. 1c). Surrounding the vessel is the space of Disse with stellate cells in a lower density collagen matrix and a concentric layer of higher density collagen with HCC cells (Fig. 1d). Confirmation of the presence of space of Disse and different cellular morphologies was detected using scanning electron microscopy (Fig.s 1e and S3). HCC cells exhibited a circular morphology with a lack of microvilli similar to *in vivo*. ECs were elongated in the direction of flow, and Kupffer macrophages presented with a more rounded morphology on the ECs, which shows liver-specific cells in HCCoC present in their native morphology.

**Figure 1:**
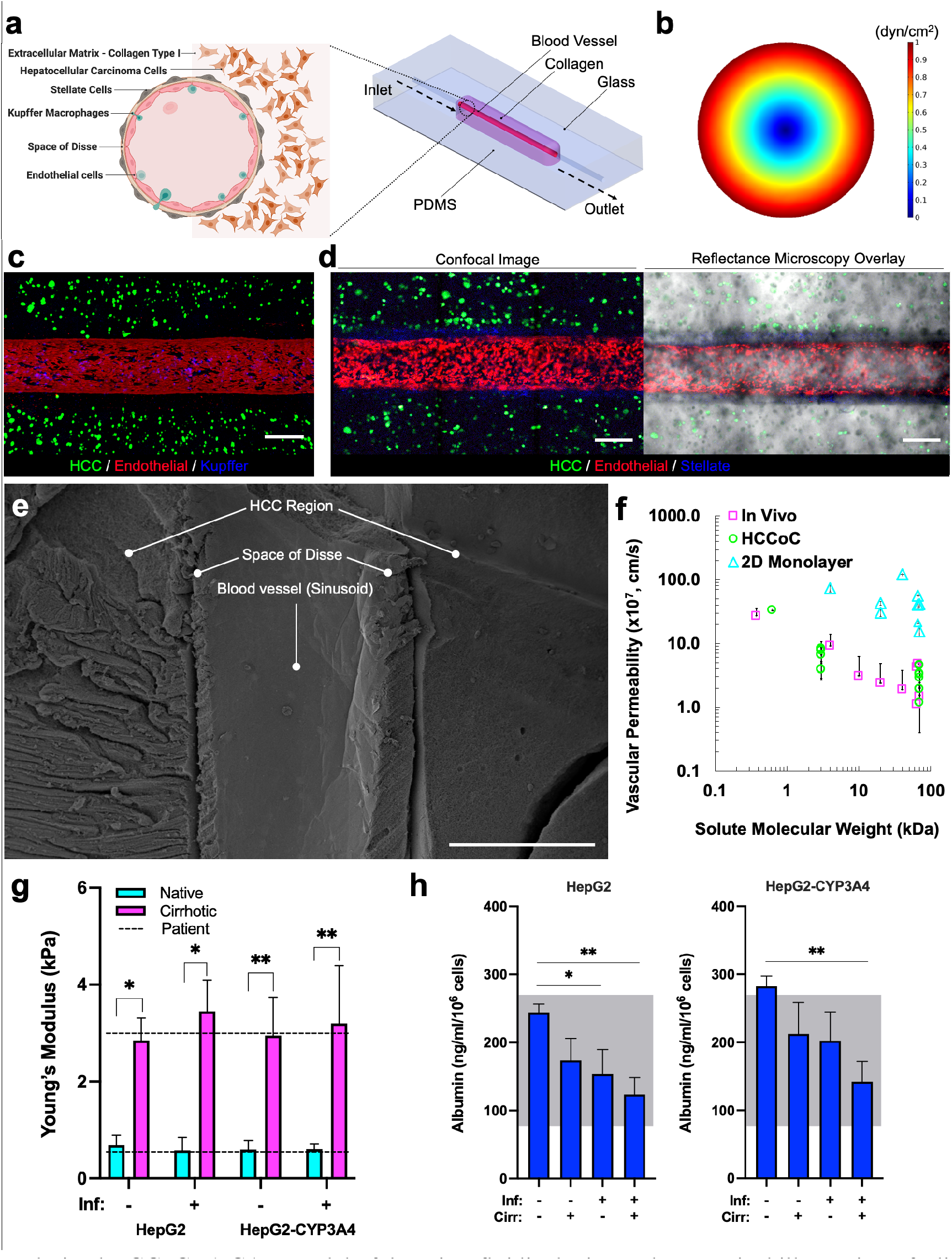
Vascularized HCCoC. **a)** CAD model of the microfluidic device and anatomical illustration of a liver sinusoid cross-section. The device consists of a single inlet and outlet with four major cell lines in liver. **b)** Finite element modeling of shear stress profile in HCCoC showing target 1 dyn/cm^2^ wall shear stress has been reached in the device. **c)** Device contains HCC (GFP), endothelial (mKate), Kupffer cells derived from THP-1 monocytes (Cell Tracker), and unlabeled stellate cells. **d)** Labeled stellate cells (Cells Tracker) in the space of Disse surrounds the circular blood vessel. **e)** SEM image of the HCCoC cross-section at the center of the blood vessel. **f)** Vascular permeability of vascularized HCC-on-a-chip, *in vivo*, and 2D monolayer findings in the literature at different solute sizes. [36–43] **g)** Compression modulus of HCCoC under normal and inflamed conditions tuned by collagen content to match native and cirrhotic stiffnesses and their comparison with HCC patient biopsy tumor reported in the literature. [44,45] **h)** Albumin secretion of HCC cells and comparison with reported HCC patient values. [46,47] All data is obtained on day three upon the completion of the preconditioning protocol. Scale: 300 μm. *p < 0.05, **p < 0.01. n = 4. All data represent means ± SD. HCC: hepatocellular carcinoma. CAD: Computer-aided design. HCCoC: Hepatocellular carcinoma-on-a-chip, GFP: green fluorescent protein. Inf.: Inflammation, Cirr: Cirrhosis.

After completion of the preconditioning protocol (3 days), the confluent endothelial barrier permeability was measured using doxorubicin (0.625 kDa) and dextran (3 kDa and 70 kDa) various *in vivo* and *in vitro* studies to measure (Fig. 1f). The vascular permeability of the HCCoC was on the order of 10^−8^ to 10^−7^ cm/s, which is comparable to *in vivo* conditions for the same tracers [36–43]. Previously reported 2D monolayer measurements demonstrates permeability values in the order of magnitude of 10^−6^ cm/s, which is significantly higher than *in vivo* studies. As a result, vascularized HCCoC device generates *in vivo*-like and representative permeability values for various dextran particle sizes.

As presented in Fig. 1g, HCCoC devices with 4 mg/mL collagen, representing native hepatic tissue, demonstrate an average stiffness of 0.69 ± 0.20 kPa and 0.60 ± 0.10 kPa for normal and inflamed states, respectively. These values are consistent with reported human liver tissue stiffness with comparable measurement techniques (0.67 ± 0.34 kPa) [44,45]. Similarly, HCCoC devices with 7 mg/mL collagen, representative of cirrhotic HCC tissue, display an average stiffness of 2.67 ± 0.62 kPa and 3.33 ± 0.92 kPa for normal and inflamed states, respectively in agreement with reported HCC patient findings (3 kPa) [44,45]. Moreover, we confirmed that the HCCoC Young’s modulus was not significantly affected by culturing different HCC cells (HepG2 or HepG2-CYP3A4) or varying THP-1 derived Kupffer cells number representing other inflammation states.

Lastly, albumin secretion in response to cirrhosis and inflammation (Fig. 1h) is comparable to measurements from HCC patients [46,47]. In agreement with previous clinical reports, we observed that implementing inflammation and cirrhosis downregulated albumin secretion in HCCoCs regardless of HCC cells [48,49]. Still, HCCoCs with HepG2-CYP3A4 cells showed higher (p<0.05) overall albumin secretion than those with HepG2 cells.

### 3.2. Cirrhotic and Inflamed HCCoCs have a compromised endothelium and elevated cytokine secretion

Liver cirrhosis and inflammation are complications of many chronic liver disease states that result in HCC [34]. They are commonly associated with HCC comorbidities that can affect vascular integrity primarily through the induction of proinflammatory cytokines (Fig. 2a) [50]. Inflammation of the liver causes the upregulation of the Kupffer cell population in the TME, which invades through the endothelial vasculature to reach HCC cells [34,35]. Activated stellate cells in the liver differentiate into myofibroblasts resulting in an upregulation of α-SMA expression, which promotes the deposition of collagen/fibrin fibers, leading to increased microenvironmental stiffness and a cirrhotic TME.

**Figure 2:**
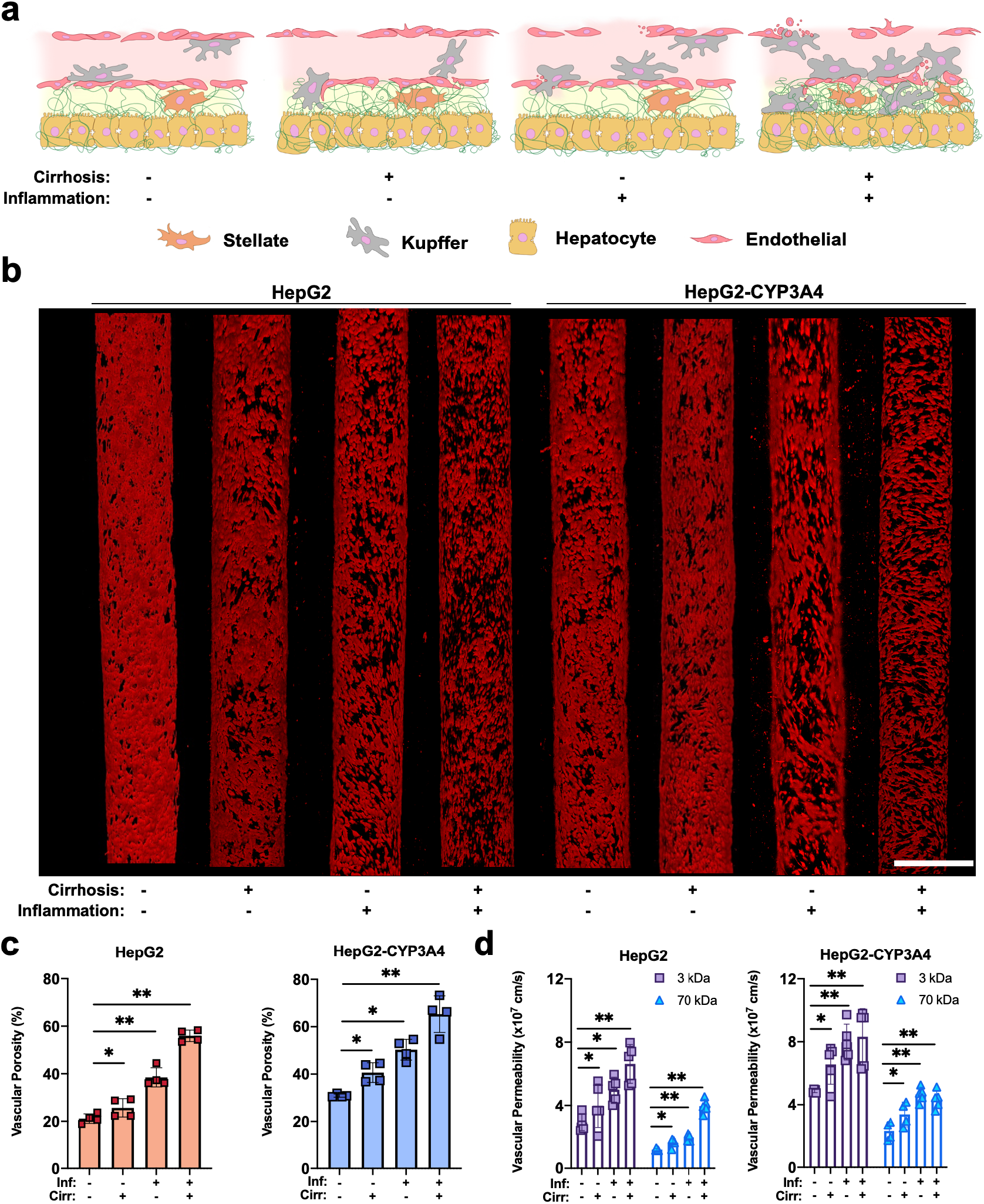
Inflammation and cirrhosis regulating the vascular and transport properties of the HCCoC. **a)** Illustration of a liver sinusoid with or without the influence of inflammation and cirrhosis. **b)** Confocal images of endothelial (red) cells under the influence of inflammation and cirrhosis in vascularized HCCoC models. Inflammation damages the vascular endothelium more than cirrhosis. Vasculature porosity was higher for the HepG2-CYP3A4 cell line (CYP3A4 overexpressing) than the low CYP3A4 expressing cell line (HepG2). **c)** Quantified vascular porosity under different microenvironmental conditions. **d)** Vascular permeability of HCCoCs with different microenvironments. An increase in vascular porosity also increased the permeability during the disease progression. Inf.: Inflammation, Cirr: Cirrhosis. HCCoC: Hepatocellular carcinoma-on-a-chip, CYP3A4: cytochrome p450-3A4 *p < 0.05, **p < 0.01. n = 4 for vascular porosity and n = 5 for permeability measurements. All data represent means ± SD. Scale is 400 μm.

Fig. 2b shows that HCCoCs without cirrhosis and inflammation have a confluent vascular endothelium with small perivascular detachments (shown as black regions in the red fluorescent vasculature). The implementation of cirrhosis in the HCCoCs increased the vascular porosities by 1.24- and 1.29-fold for HCCoCs with HepG2 and HepG2-CYP3A4 cells, respectively (Fig. 2c). The increase of ECM stiffness by tuning collagen concentration has been shown to alter EC behavior [51], suggesting that ECs in HCCoC respond to changing mechanical composition of the surrounding collagen matrix. The generation of large perivascular detachments due to the enhanced permeability retention (EPR) effect is commonly seen in tumor vessels [23]. The implementation of inflammation alone in HCCoC (represented by the increase in Kupffer cells) increased the vascular porosity by 1.86- and 1.64-fold for HCCoCs with HepG2 and HepG2-CYP3A4 cells, respectively. The combination of inflammation and cirrhosis significantly (p < 0.05) increased vascular perivascular detachments compared to the control, with an increase in vascular porosity of 2.83- and 1.90-fold for HCCoCs with HepG2 and HepG2-CYP3A4 cells, respectively. When HCCoCs cultured with HepG2 and HepG2-CYP3A4 cells are compared, the HCCoCs with the HepG2-CYP3A4 cells with upregulated CYP3A4 expression demonstrated a higher vascular porosity than HCC\oCs with the HepG2 cells in control, inflammation only, and cirrhosis only states. When inflammation and cirrhosis are combined, the vascular porosity was not significantly different (p = 0.4) between HCCoCs with HepG2 and HepG2-CYP3A4. This increase in porosity due to high CYP3A4 expression is attributed to an increased number of fenestrations present in liver sinusoid regions with upregulated metabolic gene expression *in vivo* [52].

Vascular porosity increase in response to cirrhosis and inflammation corresponds to a comparable increase in vascular permeability. Fig. 2d presents the dextran vascular permeability for HCCoC containing either HepG2 or HepG2-CYP3A4 cells with or without the influence of inflammation and cirrhosis. Paralleling HCCoC porosity data, the HCCoCs with HepG2-CYP3A4 cells present higher vascular permeability than comparable HCCoC with HepG2 cells with or without the influence of inflammation and cirrhosis alone (p < 0.05). However, vascular porosity for combined inflammation and cirrhosis conditions was similar (p = 0.1) and comparably high compared to other TME conditions. In this situation, even though the vascular porosities of HCCoCs with HepG2 and HepG2-CYP3A4 cells were similar, we were able to detect differences in vascular permeability between these two types of HCCoC using 3 kDa and 70 kDa dextran particles. This demonstrates that vascular porosity higher than approximately 70% (due to combined inflammation and cirrhosis) may not be the only indicator of vascular permeability in the HCC TME.

The regulation of vascular porosity and permeability in response to cirrhosis and inflammation could be due to the upregulation of inflammatory cytokines related to vascular leakiness and endothelium detachment such as MIP-1β, IL (interleukin)-6, IFN-γ, TNF-α, IL-1β, MCP-1, and IL-8 [50,53]. Inflammatory cytokine release increased in cirrhotic and inflammatory HCCoCs, regardless of the HepG2 subtype (Fig. 3a). Specifically, we observe a significant increase (p < 0.05) in IL-1β in inflammatory HCCoCs cultured with HepG2 and HepG2-CYP3A4 cells, which may be attributed to the increase in the macrophage population [54]. The cirrhotic condition showed minimal effect on IL-1β levels compared to control HCCoCs (without inflammation and cirrhosis). Similarly, cirrhosis increased TNF-α levels only in HCCoC with HepG2-CYP3A4 [55]. On the other hand, when cirrhosis and inflammation are combined, an increase in TNF-α levels in HCCoCs with HepG2 and HepG2-CYP3A4 was observed compared to cirrhosis alone. Macrophage inflammatory protein (MIP-1β/CCL4), produced by macrophages in response to proinflammatory cytokines, significantly increases (p < 0.05) in response to inflammation. However, unlike other measured proinflammatory molecules, we see a significant increase, but less in MIP-1β production attributed to cirrhosis potentially stemming from ECs [56]. Monocyte chemoattractant protein-1 (MCP-1/CCL2), produced mainly by macrophages and fibroblasts to help regulate macrophage migration and infiltration, shows the presence of cirrhosis or inflammation alone in the HCCoCs with HepG2-CYP3A4 [57]. However, there is a significant increase (p < 0.05) in MCP-1 production in both HCCoCs with HepG2 and HepG2-CYP3A4 in response to the combination of inflammation and cirrhosis compared to other disease states alone. This may indicate that both inflammation and cirrhosis need to be present in HCC to regulate migration and infiltration of monocytes or macrophages. Comparable to MCP-1, we see a moderate increase in interferon-gamma (IFN-γ) in response to cirrhosis (1.8-fold, p < 0.05) or inflammation (2.7-fold, p < 0.05) alone within the HCCoCs with HepG2-CYP3A4 only as shown in Fig. 3a. IFN-γ within the HCCoC could be attributed to macrophages, as limited studies have demonstrated the IFN-γ producing capability of stimulated macrophages *in vitro* [58]. The macrophage chemoattractant IL-8 is primarily produced by the non-parenchymal cells in the HCC tumor microenvironment, including activated hepatic stellate cells and increased stiffness [59]. In the HCCoCs, we see a significant increase (p < 0.01) in IL-8, regardless of which HCC cell is cultured in response to cirrhosis only, followed by a more notable increase in inflammation only, with the highest levels of IL-8 present in HCCoCs with combined inflammation and cirrhosis. IL-6 also increases proinflammatory responses in the TME, playing a role in hepatic stellate cell activation and increased stiffness [60]. In the HCCoCs, regardless of the hepatocyte cell type used, we see that inflammation alone increases IL-6 levels, likely in direct response to an increase in the macrophage population. However, the inclusion of both cirrhosis and inflammation leads to the highest production of IL-6 in all conditions tested in HepG2 (31-fold, p < 0.01) and HepG2-CYP3A4 (50-fold, p < 0.01), demonstrating a role of the microenvironment in regulating IL-6 production [61].

**Figure 3:**
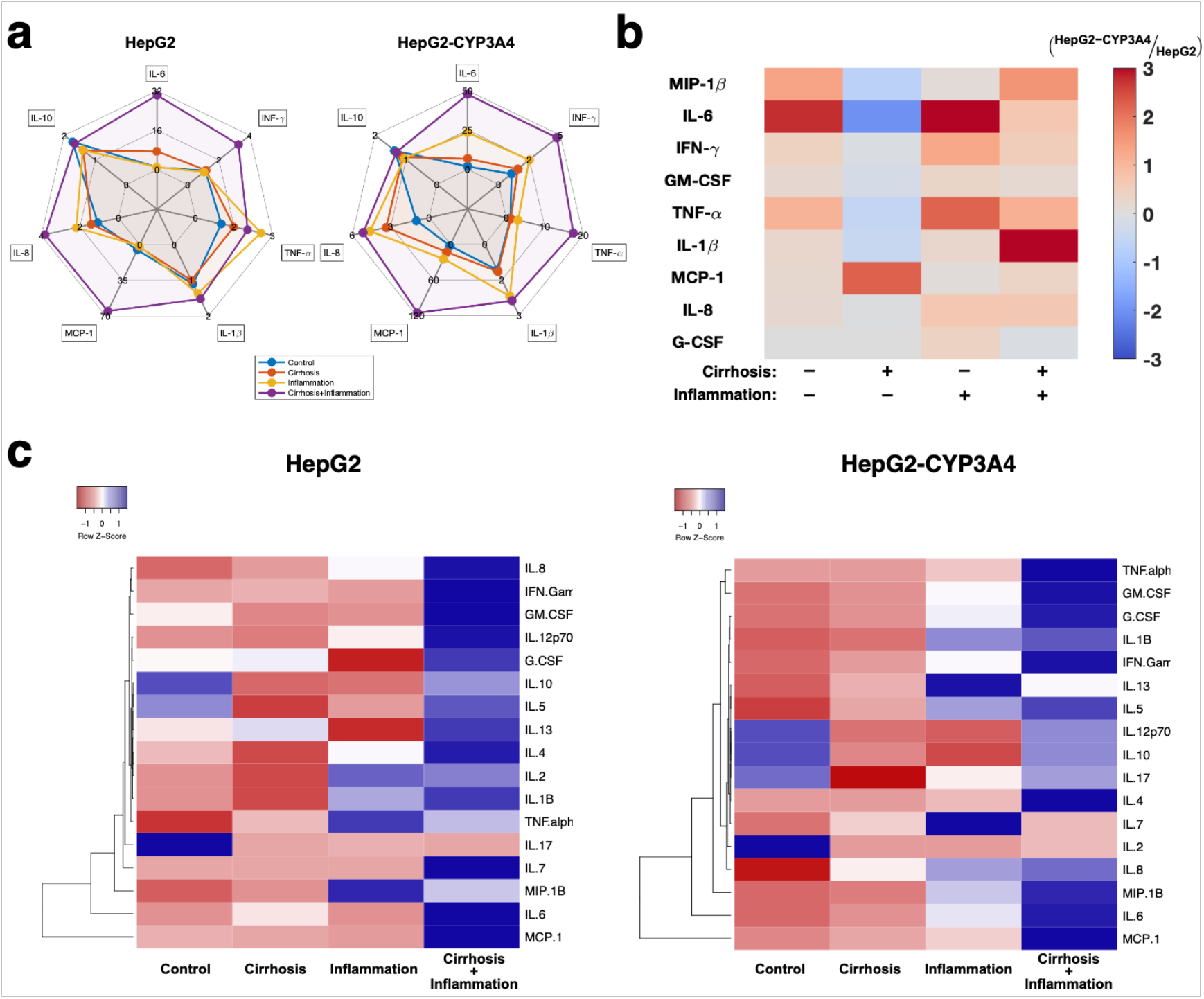
Regulation of inflammatory markers under the influence of inflammation and cirrhosis. **a)** Quantified fold change of cytokine regulations under the influence of inflammation and cirrhosis in the HCCoC. Main inflammatory cytokines and chemokines were upregulated during the TME progression. **b)** Log2 fold change comparison between HepG2 and HepG2-CYP3A4 cell lines in HCCoC under the influence of inflammation and cirrhosis. CYP3A4 overexpressing HCCoCs secreate higher inflammatory cytokines and chemokines during the disease progression. **c)** Hierarchical clustering of cytokine/chemokine responses to cirrhosis and inflammation plotted in heatmaps. n = 3. All data represent means ± SD.

We find that the upregulation of inflammatory cytokines correlated closely with the regulation of vascular porosity and permeability. Exposing ECs to inflammatory cytokines such as TNF-α, IL-1β, and IFN-γ have been previously shown to increase endothelial permeability [62]. We found that inflammation and cirrhosis alone or together had increased vascular porosity and permeability compared to the control. The HepG2-CYP3A4 HCC cells expressed higher levels of cytokines than the HepG2 cells (Fig. 3b). When only cirrhosis is implemented, most cytokines were overexpressed in HCCoCs cultured with the HepG2 cells compared to HepG2-CYP3A4 cells. However, MCP-1 is overexpressed for the HepG2-CYP3A4 cells. This may show the dominancy of MCP-1 over other cytokines in causing vascular changes since we were still able to observe higher vascular porosity and permeability for HepG2-CYP3A4 cells. This potential mechanism of vascular permeability regulation through MCP-1 expression was also shown by Stavatovic *et al*. [63]. Furthermore, hierarchical clustering analysis shows dominant inflammatory cytokines such as MCP-1, IL-6, and MIP-1β followed similar patterns in disease progression in both HCCoC with HepG2 or HepG2-CYP3A4 (Fig. 3c). Overall, inflammatory cytokines were overexpressed in the HCCoC with upregulated CYP3A4 overexpression, which resulted in higher vascular porosity and permeability.

### 3.3 HCC treatment efficacy alternation due to delivery method, inflammation, and cirrhosis

The HCC treatment efficacy has been shown in clinical studies to be regulated by cirrhosis and inflammation [64]. The potential chemoresistance regulation of HCC cells in response to disease progression has also been linked to CYP3A4 expression [14]. First, to assess the contribution of inflammation and cirrhosis to HCC chemoresistance, we investigated the treatment efficacy of doxorubicin, which is delivered either with IV or TACE (Fig. 4a).

**Figure 4:**
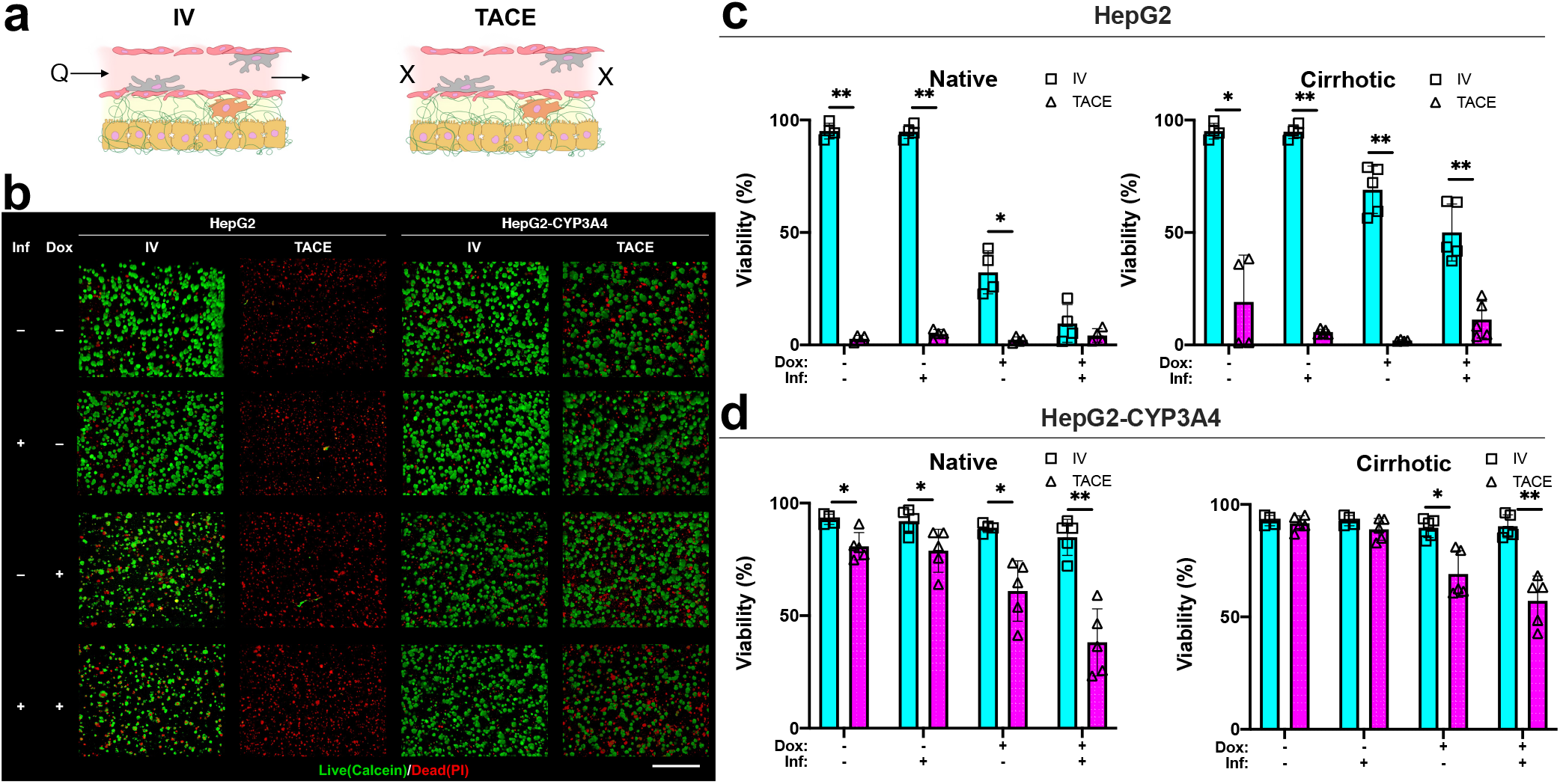
Drug treatment efficacy of IV and TACE delivery methods under the influence of inflammation and cirrhosis. **a)** Illustration of drug delivery strategy. Doxorubicin was continuously perfused in IV or embolized in TACE. Sample images refer to cirrhotic condition. **b)** Sample confocal images of live (Calcein AM, green) and dead (PI, red) HCC populations in HCCoCs. Sample images are from HCCoCs with cirrhotic stiffness and normal state. TACE improves treatement efficacy to doxorubicin. Quantified viability of **c)** HepG2 **d)** HepG2-CYP3A4 populations after the treatment with doxorubicin. Samples were compared between IV and TACE methods under the same conditions. Inflammation improves the treatment efficacy as opposite to cirrhosis, which improves the efficacy. TACE improves treatment efficacy compared to IV with under conditions of inflammation or cirrhosis. *p < 0.05, **p < 0.005. n = 4. All data represent means ± SD. Scale is 500 μm. IV: Intravenous, TACE: transcatheter arterial chemoembolization. HCCoC: Hepatocellular carcinoma-on-a-chip. Inf.: Inflammation. Dox: Doxorubicin.

Fig. 4b shows sample live and dead HCC populations stained with Calcein and PI for visualization and quantification following treatment. Quantified HCC viability 24 h after treatment with doxorubicin using IV and TACE for HepG2 (Fig. 4c) and HepG2-CYP3A4 (Fig. 4d) are shown. Our results demonstrate that delivery of doxorubicin with TACE was highly effective in HCCoCs with HepG2 cells. Even embolization without doxorubicin (Transarterial Embolization, TAE) terminated the total population of HepG2 cells in HCCoCs. However, we saw that the chemotherapeutic efficacy of TACE was lower for HCCoCs cultured with HepG2-CYP3A4 compared to HepG2 cells, which could be due to the upregulated CYP3A4 expression and increased metabolism of doxorubicin. On the other hand, doxorubicin treatment with the IV method was not as effective as TACE on HepG2-CYP3A4 cells.

In all cases, we observed that TAE was effective at terminating a proportion of the HCC tumor cell population. Blocking the incoming flow to the tumor vessel in TAE has been shown to decrease nutrient and oxygen concentration in the TME, which further causes starvation and increased HCC cells’ sensitivity to doxorubicin (Fig. S4) [65]. All tested cases in HCCoCs with the HepG2 cells show that embolization alone terminates all the HCC population without the presence of doxorubicin. In HCCoCs cultured with HepG2-CYP3A4 cells, TAE alone was able to terminate nearly 20% of HCC cells in an ECM of native stiffness, which is much less than TAE’s efficacy on HepG2 (100%). This could be the reason why the effectiveness of TACE could be influenced by CYP3A4 expression. Clinical observation of highly variable HCC CYP3A4 expression has been observed previously [13]. Potentially, a lower dose of doxorubicin could be used on the patients with lower CYP3A4 expression to reach the same drug efficacy with less off-target hepatotoxicity.

The effect of inflammation by itself did not alter the drug treatment response in native or cirrhotic TMEs. In several cases, we observed inflammation increased the efficacy of delivered doxorubicin. These cases include IV delivery in HepG2 with native (17% decrease in viability, p < 0.05) or cirrhotic (23% decrease in viability, p < 0.05) stiffnesses, TACE delivery in HepG2-CYP3A4 with native (15% decrease in viability, p < 0.05) or cirrhotic (10% decrease in viability, p < 0.05) stiffness. However, when doxorubicin is delivered with TACE to HepG2 and delivered with IV to HepG2-CYP3A4, cell viability does not change due to inflammation. This is due to the delivered doxorubicin already terminating the HCC population or was insufficient to cause toxicity. Subsequently, CYP3A4 regulation results given in Fig. 5a shows that CYP3A4 expression is downregulated when the inflammation is implemented in the HCCoC. Under the light of clinical reports, exposure of hepatic cells to TNF-α, IL-1β, IL-6, and INF-γ as a result of inflammation showed downregulation of cytochrome P450 markers [66]. In the previous section, we showed that the secretion of these cytokines in the HCCoCs was upregulated in response to inflammation. This shows HCCoC’s response to inflammation follows the same pattern observed in the patients.

**Figure 5:**
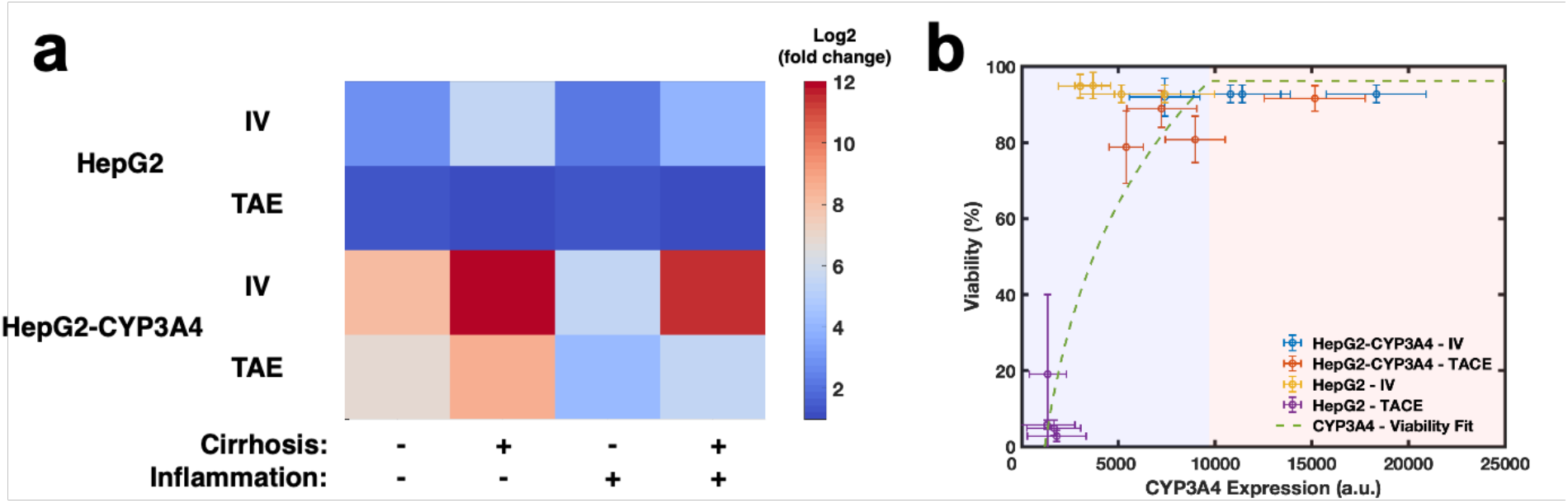
Regulation of normalized CYP3A4 expression under the influence of inflammation and cirrhosis with different treatment methods. **a)** Normalized log2 expression of CYP3A4 in correspondence to TME and drug delivery methods. Embolization through TAE method downregulated CYP3A4 expression in both HCC cells compared to IV delivery. Inflammation has been shown to decrease CYP3A4 expression and cirrhosis upregulated CYP3A4 expression of different HCC cells. **b)** Correlation of viability in correspondence with CYP3A4 expression. The logarithmic decrease in treatment efficacy was observed as CYP3A4 expression increased. Blue and red regions show effective and ineffective metabolic activity regions based on viability change, respectively. Error bars represent standard deviation. HCC: hepatocelluar carcinoma. TAE: transarterial embolization. TACE: transcatheter arterial chemoembolization. IV: Intravenous.

The stiffness increase due to cirrhosis in the TME has been shown to alter the outcome of the chemotherapeutic treatment *in vitro* [67]. In most of our findings, particularly for IV treatment for HepG2, HepG2-CYP3A4, and TACE treatments of HCCoCs with both cells, increased stiffness significantly decreased treatment efficacy. Similarly, we have previously shown that increased stiffness due to cirrhosis increases CYP3A4 expression and increases chemoresistance of HCC cells in 3D static gels [19]. We showed that in all cases, cirrhosis in the HCCoCs increases vascular porosity and increases the transport of solutes. In tested cases, our time-lapse doxorubicin transport images showed that the endothelium loses native morphology and detaches from the lumen due to the drug exposure.

Lastly, regulation of CYP3A4 expression in response to TME properties such as inflammation and cirrhosis, and drug delivery methods such as IV and TAE without the presence of doxorubicin also showed a distinguishable variation under the influence of inflammation and cirrhosis alone or combined. Overall, cirrhosis has been shown to upregulate CYP3A4 expression, and inflammation downregulated CYP3A4 expression. The alternation of CYP3A4 expression in response to TME is attributed to upregulated cytokines, such as IL-6, IL-1β, and TNF-α, in response to inflammation downregulated CYP expressions, which further diminishes drug clearance and increases drug toxicity, in HCC as well as *in vivo* HCC TMEs [14,68]. A similar trend was not observed when TACE was applied on HepG2 cells because HepG2 cells were easily affected and terminated during this process, as we have shown in Fig. 4b.

This suggests below a certain threshold of CYP3A4 expression, HCC does not demonstrate a significant chemoresistance to doxorubicin, and are subsequently terminated. In summary, Fig. 5b collectively shows an increase in CYP3A4 expression of HCC cells and a corresponding increase in their viability in response to treatment. Overall, the regulation of CYP3A4 parallels what we observed in drug treatment results, which suggests CYP3A4 expression could be a predictive treatment outcome in HCC. Furthermore, the observed difference of efficacy between HCCoCs with the two cell types mirrors clinical observation of variable HCC CYP3A4 expression [13].

### 3.4. TACE delivered doxorubicin induces less vascular damage than IV delivery

Despite the advantage of doxorubicin in terminating HCC populations, non-targeted cytotoxic damage has been one of the greatest drawbacks. As ECs regulate drug flow from the vasculature to the hepatic region, they are the first cell type exposed to doxorubicin. Dysfunction of the endothelium would further impact the regulation of macromolecule transport from the vasculature to the hepatic region.

In this section, we quantify how delivering doxorubicin using TACE or IV affects vascular porosity. Fig. 6 shows vascular porosity quantified after doxorubicin treatment in response to inflammation and/or cirrhosis. Confocal images shown in Fig. 6a show vascular integrity after TACE or IV treatment with or without delivering the doxorubicin. Treatment within vascularized HCCoC showed that perivascular detachment occurs due to chemotherapy regardless of the delivery method. The IV-delivered doxorubicin created a dysfunctional damaged endothelium with the contribution of shearing off damaged ECs. However, TACE delivered doxorubicin did not increase vascular damage as much as IV delivery. Time-lapse doxorubicin transport images also revealed that applied WSS due to continuous flow contributes to perivascular detachment of ECs together with doxorubicin during IV treatment. TACE without doxorubicin also increased perivascular detachment, but did not result in a severely non-functional endothelium. The vascular porosity after TACE or IV combined with doxorubicin for HCCoC containing HepG2 (Fig. 6b) and HepG2-CYP3A4 cells (Fig. 6c) with or without the influence of inflammation and cirrhosis alone or combined were quantified. Regardless of the presence of inflammation or cirrhosis, IV-delivered doxorubicin resulted in a significantly higher increase in vascular porosity than TACE delivery.Our findings show an increase in vascular porosity was attributed to the presence of doxorubicin. In the light of preclinical studies [69], the increase in porosity possibly increased the vascular permeability of tumor vessels.

**Figure 6:**
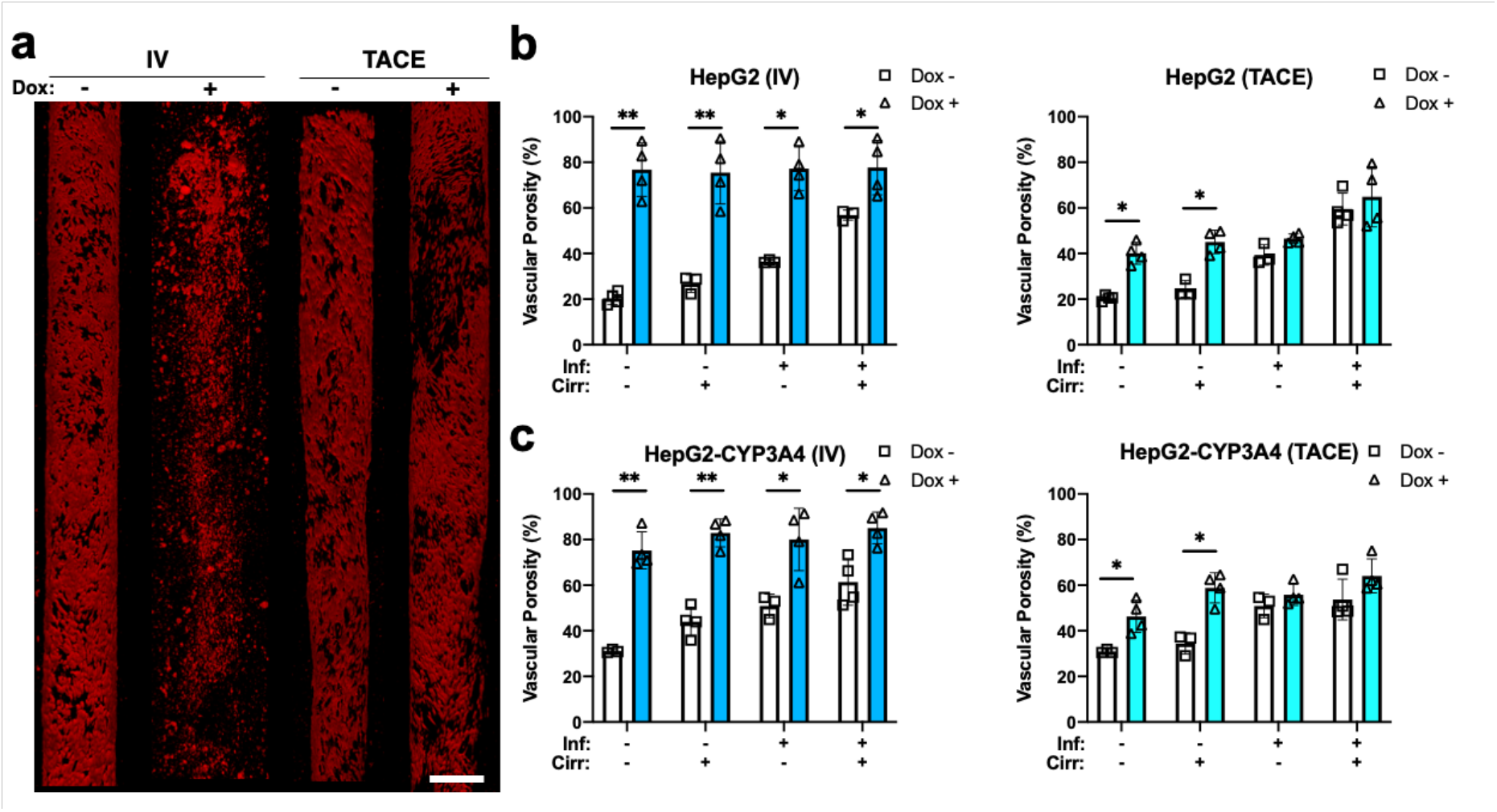
Vascular damage created by doxorubicin treatment. **a)** Sample vessel images of endothelial (red) treated with or without doxorubicin using TACE and IV methods. Sample images are from HepG2 HCCoCs with native stiffness and normal state. Quantified vascular porosity change with and without the influence of doxorubicin delivered by IV and TACE methods on **b)** HepG2 and **c)** HepG2-CYP3A4 HCC cells. Nonselective vascular porosity increases 24 h after 10 μM doxorubicin was delivered using IV or TACE methods. *p < 0.05, **p < 0.01. n = 4. All data represent means ± SD. Scale is 300 μm. IV: Intravenous. TACE: transcatheter arterial chemoembolization. Inf.: Inflammation, Cirr: Cirrhosis, Dox: Doxorubicin

## 4. Discussion and Conclusions

The measurement of drug and solute transport across the endothelium is a critical, unmet need for assessing the potential efficacy of biotherapeutics and alternative delivery methods under different states of HCC progression. We have created the first vascularized HCCoC model to replicate functional albumin secretion and *in vivo*-like vascular permeability for measuring transport. HCCoCs’ tunable tumor microenvironment properties such as ECM Young’s modulus to simulate cirrhosis and local Kupffer macrophage population to reproduce various inflammation states allow further investigation of how disease progression alters vascular properties and HCC chemoresistance. Furthermore, HCCoC enables the characterization of tissue response to multiple drug delivery methods.

In this work, we quantified how vascular porosity and permeability could be modulated by inducing inflammation and cirrhosis disease states, which contribute to the upregulation of inflammatory cytokines, primarily by MCP-1, IL-6, and MIP-1β. This upregulation and subsequent modulation of vascular porosity reuluts solute transport is driven by paracellular transport rather than transendothelial flux in HCC [70]. Furthermore, the HCCoC provided a high resolution, spatiotemporal investigation of the efficacy of chemotherapeutics on tumor cells and vasculature.

Our findings show that cirrhosis, inflammation, and CYP3A4 metabolic activity increase vascular permeability due to the upregulation of inflammatory cytokines. Between cirrhosis and inflammation disease states, cirrhosis induced inflammatory cytokines similar to inflammation but at lower levels. Furthermore, high CYP3A4 expression in HCC cells can be a key factor in regulating vascular properties through upregulation of cytokines responsible for perivascular detachment through secretion of TNF-α, IL-6, and MCP-1 [71]. As a result of the link between CYP3A4 and cytokine secretion, we can conclude that the vascular properties rely on metabolic activity in the TME. These findings of increased vascular permeability and porosity with CYP3A4 expression *in vitro* correlate with *in vivo* observations of increased liver fenestration (endothelial porosity) along the increasing cytochrome P450 family activity gradient from the portal vein to the hepatic artery [52].

CYP3A4 metabolic activity was also found to regulate the chemoresistance of HCC cells in response to both IV and TACE treatment methods. In general, this work shows that CYP3A4 expression was upregulated by cirrhosis and downregulated by inflammation, which is reflected in chemoresistance alteration. Overall, CYP3A4 has been shown to regulate chemotherapeutic efficacy of doxorubicin, which shows CYP3A4 expression can be a good candidate in the assessment of required dose of therapeutics and effectiveness in patients [20]. Our study further showed delivery of therapeutics through TACE downregulates CYP3A4 expression and results in higher efficacy of doxorubicin, which highlights the critical role that embolization delivery methods play in HCC treatment. This phenomenon is potentially attributed to an increase in VEGF, and oxygen depletion due to the embolization process, resulting in the regulation of CYP3A4 expression [19]. This could also increase the production of reactive oxygen species (ROS) and decrease multidrug resistance-1 (MDR-1) expression that could diminish the chemoresistance of HCC cells. In the absence of chemotherapeutic drugs, the process of embolization alone was quantitatively shown to terminate a significant population of HCC cells both in this study and clinical practices [72]. Alternatively, transient gene therapies or inhibitors to temporarily downregulate CYP3A4 expression in HCC tumors could improve treatment efficacy and be investigated in future studies. Several inhibitors include, but are not limited to, clarithromycin, diltiazem, erythromycin, itraconazole, ketoconazole, ritonavir, and verapamil. Furthermore, ECs still retain the physical confluence after TACE delivery, suggesting TACE has lower non-specific damage to ECs than IV delivery. The EC deficiency and detachment could result from shearing and drug-induced apoptosis.

In summary, this study presents a microfluidic HCCoC to investigate how the TME and drug delivery methods can regulate HCC chemoresistance, vascular permeability, and solute transport. Additionally, the need for small amounts of reagents and cells provides an inexpensive alternative to *in vivo* models. The capability to measure the spatial and temporal response in a controlled microenvironment can permit quantitative assessment of vascular, therapeutic, and gene expressions to be measured that are impossible *in vivo*. This HCCoC also provides a low-cost option for drug screening and discovery and high throughput quantification of novel HCC treatments, which could not be tested due to the limitation of technology and tunability up to this point. The potential to incorporate clinical samples in the HCCoC, can permit our microfluidic model to be translated to personalized medicine. The ability of this HCCoC to model disease progression also offers the possibility to study other patient-specific physiological and pathological processes in HCC, and further embolization strategies.

## Supporting Information

Supporting Information is available from the Wiley Online Library or from the author.

## Acknowledgements

The authors acknowledge the support of the Cancer Prevention Research Institute of Texas (CPRIT) for funding part of this work through grant RR160005, National Cancer Institutes for funding through R21EB019646, R01CA186193, R01CA201127-01A1, U24CA226110 and U01CA174706, National Institute of Diabetes and Digestive and Kidney Diseases funding through awards R01DK082435 and R01DK112803 and Department of Veterans Affairs Biomedical Laboratory Research and Development Service funding through award BX003486. The authors would like to thank Dr. Wei Li (Department of Mechanical Engineering, the University of Texas at Austin) for generously gifting CYP3A4 overexpressing cell type. T.E.Y. is a CPRIT scholar in cancer research. This material is the result of work supported with resources and the use of facilities at the Central Texas Veterans Health Care System, Temple, Texas. The content is the responsibility of the author(s) alone and does not necessarily reflect the views or policies of the Department of Veterans Affairs or the United States Government.

## Author contributions

Conceptualization: A.Ö. conceived of the idea for the study. Supervision: E.N.K.C, T.E.Y. and M.N.R. supervised the project. E.N.K.C., M.M., T.E.Y. and M.N.R provided feedback and assistance with manuscript preparation. Investigation: A.Ö., D.L.S., and M.N.R were responsible for performing the studies and analyzing the experimental data. Writing: A.Ö. wrote the initial draft of the paper. All authors discussed the results and revised the manuscript.

