## Supplementary Information for "Cirrhosis and Inflammation Regulates CYP3A4 Mediated Chemoresistance in Vascularized Hepatocellular Carcinoma-on-a-chip"

### Supplementary – I: Modeling of fluid flow and oxygen concentration in HCCoC

Accordingly, flow simulations were conducted on Comsol Multiphysics software. Stoke's Law was used to quantify the flow inside the vasculature. It is assumed that flowing fluid has constant viscosity and incompressible and Reynold's number inside the vasculature is low so that we can eliminate the inertial terms.

$$\rho(\frac{\partial u}{\partial t} + u_r\frac{\partial u}{\partial r} + \frac{u_\theta}{r}\frac{\partial u}{\partial \theta} + u\frac{\partial u}{\partial x}) = -\frac{\partial P}{\partial x} + \rho g_x + \mu(\frac{1}{r}\frac{\partial}{\partial r}\left(r\frac{\partial u}{\partial r}\right) + \frac{1}{r^2}\frac{\partial^2 u}{\partial \theta^2} + \frac{\partial^2 u}{\partial x^2}) \quad (1)$$

$$u_r = \frac{\kappa \nabla P}{\zeta \mu} \quad (2)$$

$$u_x = \frac{\kappa \nabla P}{\zeta \mu} \quad (3)$$

where is $\nabla$ gradient operator and $\nabla^2$ is the square of vector Laplacian, P is pressure, u is velocity, μ is viscosity, and ρ is the density of the fluid. Simulations were run with the following conditions: constant flow rate of 59 µl/min at the inlet and outlet boundary condition of zero gauge pressure. Vessel diameter for HCCoC is set as 435 µm respectively. Additionally, flow in extracellular matrix was modelled using Darcy's Law with isotropic tissue mechanical property. ECM properties used in the simulation were porosity of the collagen of 0.49, and hydraulic permeability of the collagen of $10\text{x}10^{-15}$ ($m^2/s$). Resulting flow profile was used to calculate wall shear stress at the vessel walls. Wall shear stress of a Newtonian fluid is defined as:

$$\tau = \mu\dot{\gamma} \quad (4)$$

where, $\tau$ is shear stress, μ is fluid viscosity, and $\dot{\gamma}$ is shear rate resulting from the flow simulations. The shear stress value was displayed across the vessel cross-section. Computational domain is given in Figure #a. Endothelial layer thickness was set as 15 µm and Space of disse thickness was set as 138 µm. Shear stress profile presented in Figure #b shows targeted 1 dyn/$cm^2$ wall shear stress was reached under given flow rate and experimental conditions.

Accordingly, time dependent convective diffusion equations in transport of diluted specifies module were solved temporally over the collagen hydrogel (Equation 5) and culture medium (Equation 6):

$$\epsilon_p\frac{\partial c}{\partial t} - \nabla.(D_e\nabla c) = R \quad (5)$$

$$\frac{\partial c}{\partial t} - \nabla.(D_{media}\nabla c) = 0 \quad (6)$$

where c is the oxygen concentration, $\epsilon_p$ is porosity of the collagen hydrogel, $\rho$ is the density of media, heat capacity of media, $D_e$ is effective oxygen diffusivity, R is oxygen consumption rate by HCC cells. The diffusivity of collagen hydrogels was adjusted using Bruggemen model, where the porosity of collagen hydrogels was required to be implemented. Accordingly, Equation 7 was incorporated to the computational model:

$$D_e = \epsilon_P^{4/3}D_{collagen} \quad (7)$$

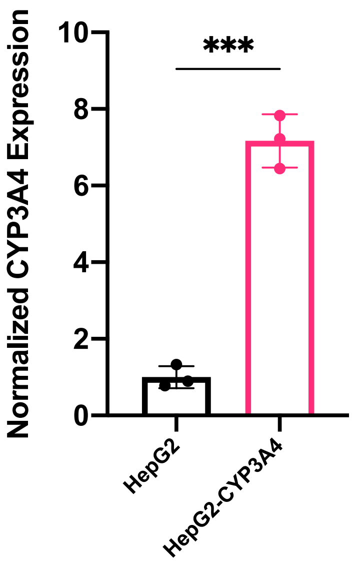


**Figure S1:** CYP3A4 expression baseline in tested HCC cell lines. CYP3A4 transfected HepG2-CYP3A4 cell lines express significantly higher metabolic enzyme compared to wild-type HepG2 cell line.

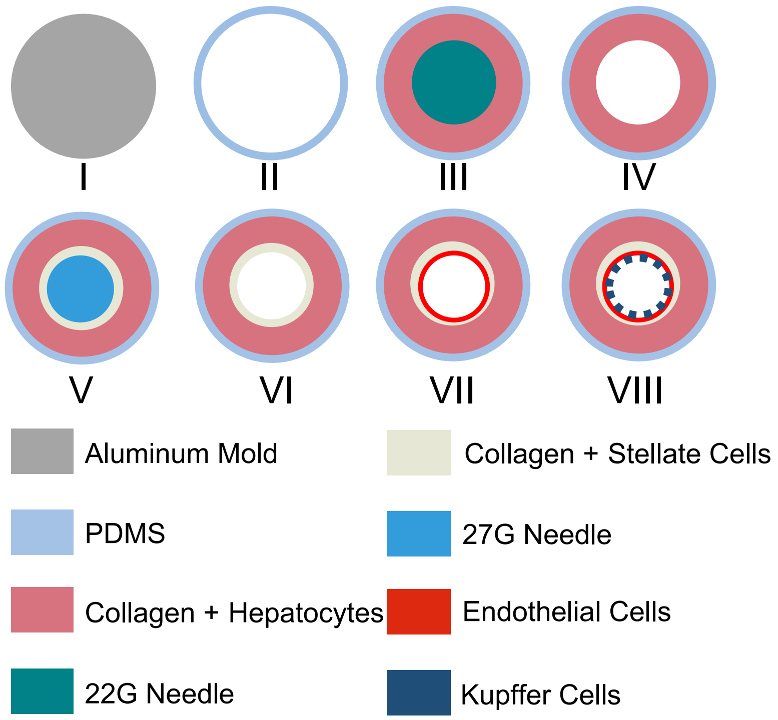


**Figure S2:** Fabrication steps of the HCCoC. HCCoC was fabricated as given in following steps. Aluminum mold shown in Figure S5-I was fabricated using micromilling technique. 22 G needle was inserted to the aluminum mold during the baking step (Figure S2-II) of the PDMS. Mixture of PDMS curing agent at 10:1 ratio was poured inside the aluminum mold and baked at 75°C for 1 hour. After this step, 22 G needle and cured PDMS were separated from the aluminum mold and PDMS was plasma bonded with glass slide under 30W exposure for 3 minutes (Figure S2-II). Collagen solution at intended concentration and HCC cell line was prepared as described in the Methods section and injected through the channels. Before polymerization of the collagen, 22G needle was inserted through the inlet of the channel to form hollow lumen (Figure S2-III). After the polymerization, needle was removed (Figure S2-IV) and second collagen solution with stellate cells were injected to the channel. Before the polymerization, 27G needle was inserted to the inlet to for a hollow lumen surrounded by thin stellate cells with collagen to form space of Disse. After the polymerization 27G need was removed (Figure S2-VI) and went under the preconditioning protocol given in Methods section to form confluent and aligned endothelium with Kupffer macrophages (Figure S2-VII and VIII).

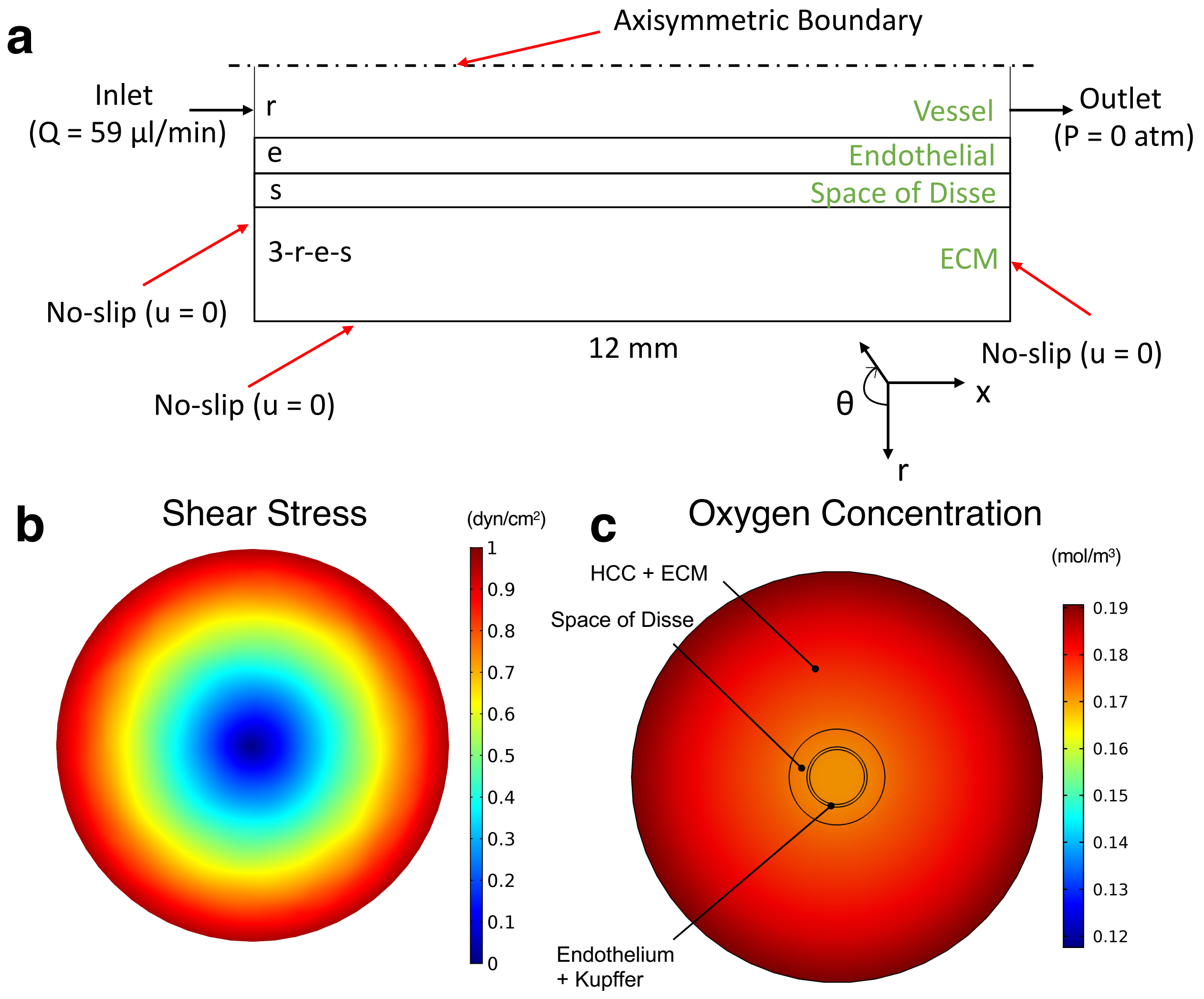


**Figure S3:** Computational modeling of flow in HCCoC. **a)** Computational domain. **b)** Shear stress profile across the vessel. Target 1 dyn/cm2 wall shear stress was reached under given experimental conditions. **c)** Spatial oxygen concentration distribution in HCCoC.

### Supplementary – II: SEM imaging of HCCoC

Aldehyde mixture composed of 0.2 M cacodylate buffer, glutaraldehyde, paraformaldehyde, cation stock, and DI $H_2O$ were prepared and fixed at room temperature for 4 hours and washed three times with cacodylate buffer for 15 minutes. Subsequently, reduced osmium solution (1:1) composed of 4% potassium ferrocyanide in 0.2 M cacodylate buffer and 4% aqueous osmium tetroxide was added to samples and maintained on ice for 4 hours. Fixed samples were washed with DI $H_2O$ 5 times for 10 minutes afterwards and dehydrated with 50, 70, and 95% ethanol once and twice with 100% ethanol for 15 minutes each. Samples were dried using a critical point drying method, coated with 12$\mu$m thick Pt/Pd layer and imaged with Zeiss Supra40 SEM. All reagents were purchased though Electron Microscopy Sciences, PA.

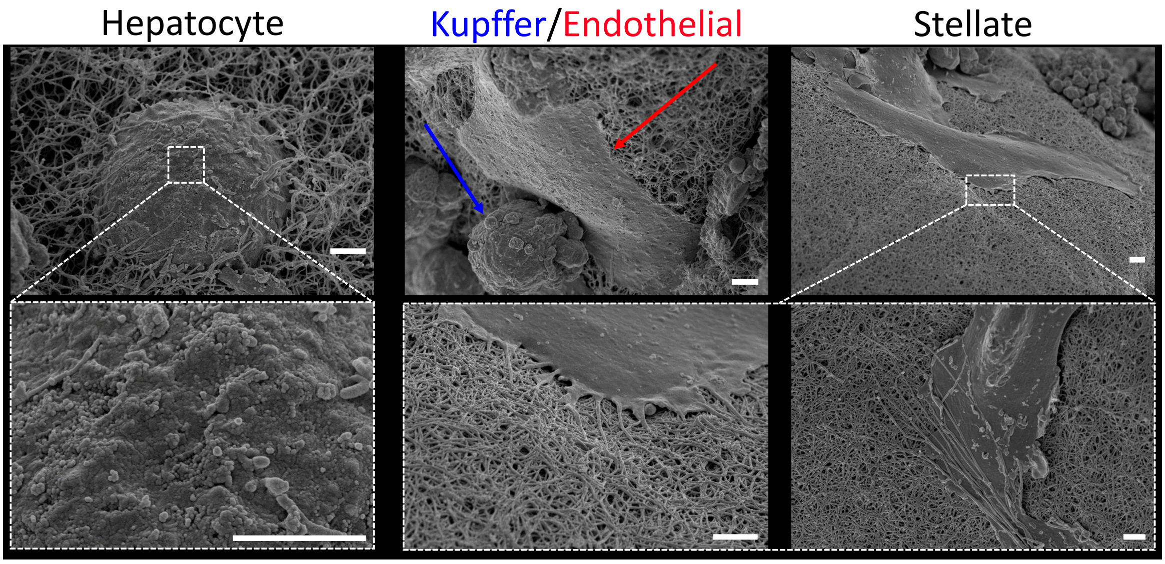


**Figure S3:** Morphology of four major cells in liver. Abundant microvilli on hepatocyte cell surface has been observed. Endothelial and Kupffer cells are interacting on the liver sinusoid. Stellate cells are blending with the collagen in the space of Disse. Scales are 2 $\mu$m.

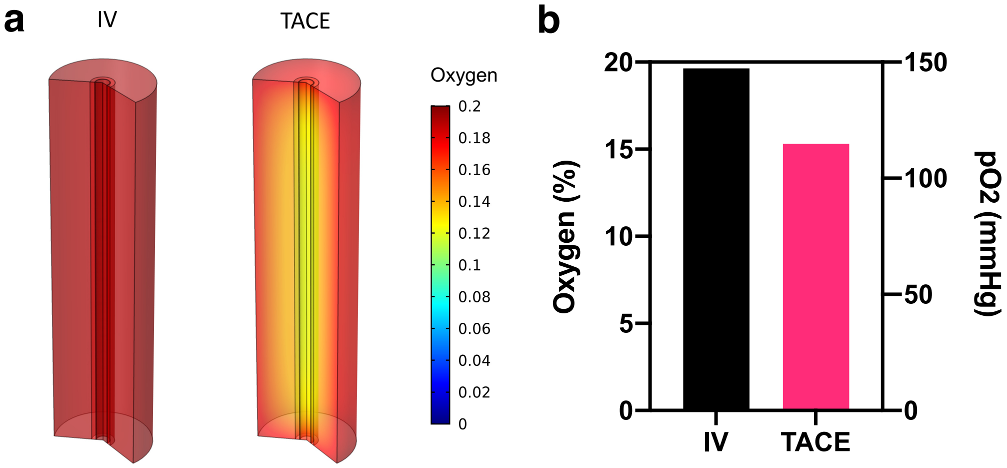


**Figure S4:** Oxygen depletion in HCCoC as a result of IV or TACE drug delivery methods. **a)** Contours of oxygen profiles in the overall device. **b)** Quantified oxygen levels in the region of HCC cells. TACE results in oxygen depletion in HCCoC.
